# Increases in multiple resources promote plant invasion

**DOI:** 10.1101/2021.08.12.456056

**Authors:** Zhijie Zhang, Yanjie Liu, Angelina Hardrath, Huifei Jin, Mark van Kleunen

## Abstract

Invasion by alien plants is frequently attributed to increased resource availabilities. Still, our understanding is mainly based on effects of single resources despite the fact that plants rely on multiple resources. How multiple resources affect success of alien plants remains largely unexplored. Here, with two common garden experiments, one in China and one in Germany, we tested whether nutrient and light availabilities affected the competitive outcomes between alien and native plants. We found that under low resource availabilities or with addition of only one type of resource aliens were not more competitive than natives. However, with a joint increase of nutrients and light intensity, aliens outcompeted natives. Our finding indicates that addition of multiple resources could greatly reduce the number of limiting factors (i.e. niche dimensionality), and that this favors the dominance of alien species. It also indicates that habitats experiencing multiple global changes might be more vulnerable to plant invasion.

## Introduction

The rapid accumulation of alien species is one of the characteristics of the Anthropocene (Lewis & Maslin 2015; van Kleunen *et al*. 2015). Because some alien species can threaten native species and disrupt ecosystem functioning (Vilà *et al*. 2011), it has become an urgent quest to understand the mechanisms that allows aliens to outcompete natives. One widely considered mechanism is proposed by the fluctuating resource hypothesis (Davis *et al*. 2000), which poses that ‘a plant community becomes more susceptible to invasion whenever there is an increase in the amount of unused resources’. Numerous studies have shown that resource increases can favor alien plants over natives (Parepa *et al*. 2013; Liu & van Kleunen 2017). However, while plants need different types of resources (e.g. nutrients, light), most studies investigated only one single resource, mainly nutrients. The few studies that investigated the effects of multiple resources on alien and native plants (Burns 2004; Dawson *et al*. 2012), usually looked at growth of individually grown plants, instead of at competitive outcomes. Therefore, it remains largely unknown whether multiple resources interact in their effects on competitive outcomes between alien and native plants.

How resources affect competition has long fascinated and puzzled ecologists (Hutchinson 1961; Chesson 2000). Resource-competition theory predicts that if multiple species are competing for resources, coexistence of all species is only possible when each species is limited by a different resource (Tilman 1982). A classic example comes from algae, where *Asterionella formosa* and *Cyclotella meneghiniana* are able to coexist when *A. formosa* is limited by phosphate and *C. meneghiniana* is limited by silicate (Tilman 1977). Resource addition (e.g. phosphate) will decrease the number of limiting resources and will thus favor dominance of one species (known as the niche-dimension hypothesis *sensu* Harpole & Tilman 2007; Fig. 1). Although it remains challenging to identify limiting resources for more complex species (e.g. vascular plants), a few follow-up experiments have shown that coexistence of multiple plant species is less likely with addition of multiple resources (Harpole & Tilman 2007; Harpole *et al*. 2016). The explanation behind this is straightforward: the more types of resources are added, the more likely it is that the previously limiting resources are no longer limiting. A next step in this field of research is to predict which type of species will be favored with addition of multiple resources.

**Figure 1.**
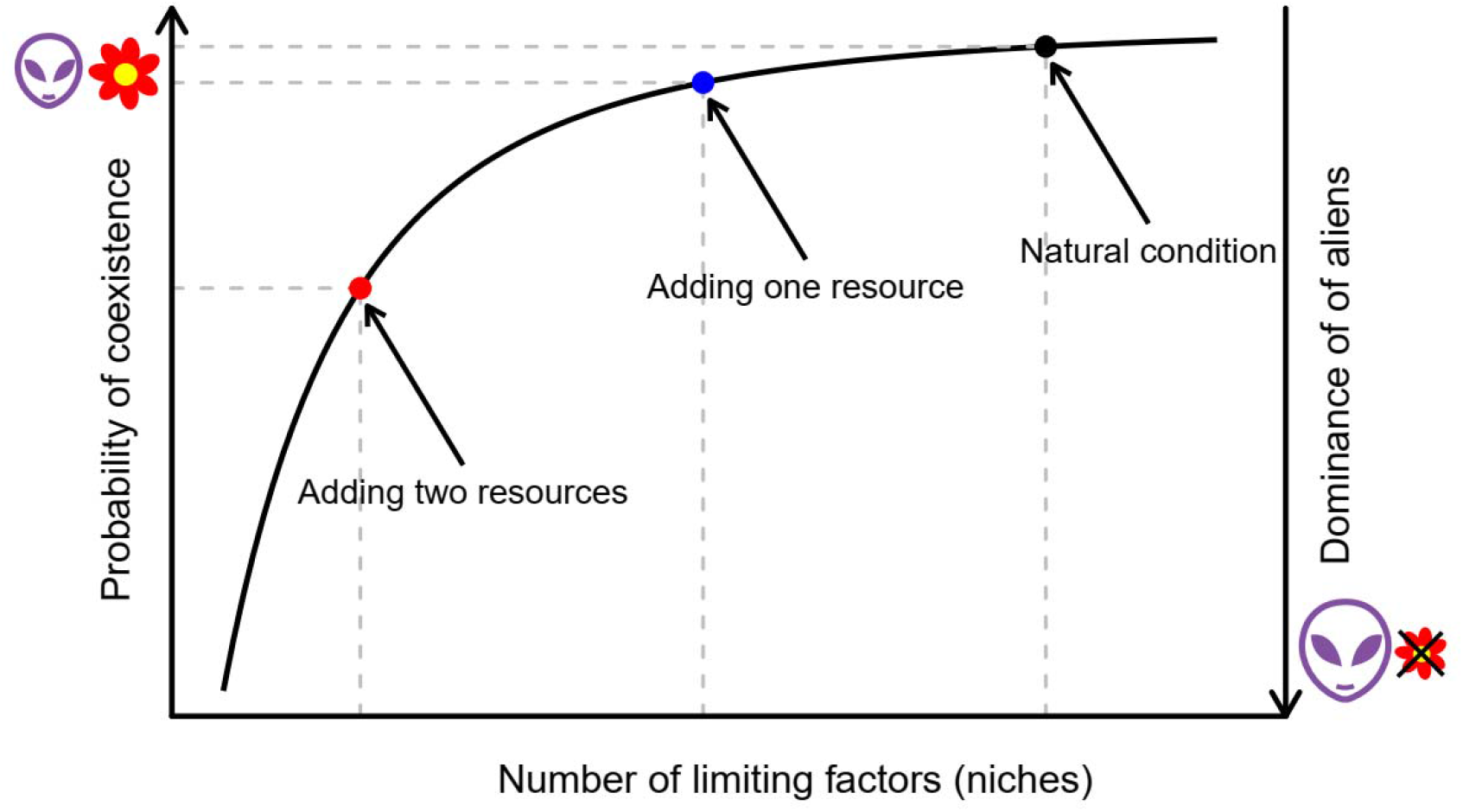
The effect of number of limiting factors on species coexistence (the niche dimension hypothesis) and its application in invasion ecology. Probability of species coexistence increases with number of limiting factors, asymptotically approaching 100%. This is because the higher number of limiting factors, the more likely that different species are constrained by different limiting factors. In natural condition, there are multiple limiting factors, such as nutrients, light and enemies, favoring species coexistence (black dot). Adding one resource will eliminate one limiting factor, slightly reducing the probability of coexistence (blue dot). Adding multiple resources will eliminate more limiting factors, greatly reducing the probability of coexistence and favoring dominance of single species (red dot). Because alien species are likely to pre-adapt to high resource availabilities and/or to be limited by fewer factors, their dominance is expected to be favored by addition of multiple resources.

One group of species that might benefit from addition of multiple resources are alien species. First, most alien plants originate from anthropogenic habitats (Kalusová *et al*. 2017), which are frequently rich in resources due to disturbance. Consequently, successful aliens are frequently those that are pre-adapted to high resource availabilities (Dostál *et al*. 2013) and thus are favored by resource addition (Seabloom *et al*. 2015; Liu & van Kleunen 2017). Second, aliens might be limited by fewer factors than natives are, because their evolutionary history differs from the one of natives (Darwin 1859; Saul *et al*. 2013). For example, alien plants might be released from and thus be less limited by natural enemies (Keane & Crawley 2002). Such advantage of aliens over natives may be expressed when both types of species suffer from resource limitation, especially from the limitation of multiple resources. However, the advantage will appear when resource limitation is removed by resource addition. Although resource-competition theory offers a potential mechanistic explanation of the success of alien species, empirical tests remain rare.

Here, we conducted two experiments, one in Germany and one in China, with similar designs. In both locations, we grew multiple alien and native plant species either alone, in monoculture, or in mixture of two species. To vary resource availabilities, we used two levels of nutrients, and two levels of light. We focused on nutrients and light for two main reasons. Frist, most successful alien plants origin from, and also have naturalized in, nutrient-rich habitats (Kalusová *et al*. 2017; Stotz *et al*. 2019), which indicates the importance of nutrients for alien plant invasion. Second, alien plants are frequently larger than native plants (van Kleunen *et al*. 2010), which might provide them with a superior ability to capture light and to throw shade on native plants. We aimed to test whether resource availabilities affected pairwise competitive outcomes between alien and native species. We expected that 1) an increase in a resource favors aliens over natives, and 2) that this effect is stronger when two resources are increased instead of just one.

## Methods and Materials

### Study species

To increase our ability to generalize the results, we conducted multispecies experiments (van Kleunen *et al*. 2014). The experiments were designed independently, but, as they used similar treatments, we analyzed them jointly to further increase generalizability. For the experiment in China, we selected eight species that are either native or alien in China (Table S1). For the experiment in Germany, we selected 16 species that are either native or alien in Germany (Table S1). All 24 species, representing seven families, are common in their respective regions. To control for phylogenetic non-independence of species, we selected at least one alien and one native species in each of the seven families. All alien species are naturalized (*sensu* Richardson *et al*. 2000) in the country where the respective experiment was conducted. We classified the species as naturalized alien or native to China or Germany based on the following databases: (1) “The Checklist of the Alien Invasive Plants in China” (Ma & Li 2018), (2) the Flora of China (www.efloras.org) and (3) BiolFlor (www.ufz.de/biolflor). Seeds or stem fragments of the study species were obtained from botanical gardens, commercial seed companies, or from wild populations (Table S1).

### Experimental set-up

#### The experiment in China

From 21 May to 27 June 2020, we planted or sowed the eight study species into plastic trays filled with potting soil (Pindstrup Plus, Pindstrup Mosebrug A/S, Denmark). To ensure that the species were in similar developmental stages at the beginning of the experiment, we sowed the species at different times (Table S1). Three species were grown from stem fragments because they mainly rely on clonal propagation, and the others were propagated from seeds (Table S1).

On 13 July 2020, we transplanted the cuttings or seedlings into 2.5-L circular plastic pots filled with a mixture of sand and vermiculite (1:1 v/v). Three competition treatments were imposed: 1) competition-free, in which plants were grown alone; 2) intraspecific competition, in which two individuals of the same species were grown together; 3) interspecific competition, in which two individuals, each from a different species were grown together. We grew all eight species without competition, in intraspecific competition and in all 28 possible pairs of interspecific competition. For the competition-free and intraspecific-competition treatments, we replicated each species seven times. For the interspecific-competition treatment, for which we had many pairs of species, we replicated each pair two times.

The experiment took place in a greenhouse at the Northeast Institute of Geography and Agroecology, Chinese Academy of Sciences (Changchun, China). The greenhouse had transparent plastic film on the top, which reduced the ambient light intensity by 12%. It was open on the side, so that insects and other organisms could enter. To vary nutrient availabilities, we applied to each pot either 5g (low nutrient treatment) or 10g (high nutrient treatment) of a slow-release fertilizer (Osmocote® Exact Standard, Everris International B.V., Geldermalsen, The Netherlands; 15% N + 9% P_2_O_5_ + 12% K_2_O + 2% MgO + trace elements). To vary light availabilities, we used two cages (size: 9 m × 4.05 m × 1.8 m). One of them was covered with two layers of black netting material, which reduced the light intensity by 71% (low light-intensity treatment). The other was left uncovered (high light-intensity treatment).

The experiment totaled 672 pots ([8 no-competition × 7 replicates + 8 intraspecific-competition × 7 replicates + 28 interspecific-competition × 2 replicates] × 2 nutrient treatments × 2 light treatments). The pots were randomly assigned to positions, and were randomized once on 15 August within each block (i.e. the low or high light-intensity treatment). We watered the plants daily to avoid water limitation. On 1 September 2020, we harvested aboveground biomass of all plants. The biomass was dried at 65□ for 72h to constant weight, and then weighed to the nearest 1mg.

#### The experiment in Germany

On 15 June 2020, we sowed seeds of the 16 species into plastic trays filled with potting soil (Topferde, Einheitserde Co). On 6 July 2020, we transplanted the seedlings into 1.5-L pots filled with a mixture of potting soil and sand (1:1 v/v). Like the experiment in China, we imposed three competition treatments: competition-free, intraspecific competition and interspecific competition. However, in this experiment, which had two times more species than the experiment in China, we only included 24 randomly chosen species pairs for the interspecific-competition treatment, and all of these pairs consisted of one alien and one native species. For the competition-free treatment, we replicated each species two times. For the competition treatments, we did not use replicates for any of the species combinations, as replication of the competition treatments was provided by the large number of species pairs.

The experiment took place outdoors in the Botanical Garden of the University of Konstanz (Konstanz, Germany). To vary nutrient availabilities, we applied to each pot once a week either 100 ml of a low-concentration liquid fertilizer (low-nutrient treatment; 0.5‰ Universol ® Blue oxide fertilizer, 18% N + 11% P + 18% K + 2.5% MgO + trace elements) or 100 ml of a high-concentration of the same liquid fertilizer (high nutrient treatment; 1‰). To vary light availabilities, we used eight metal wire cages (size: 2 m × 2 m × 2 m). Four of the cages were covered with one layer of white and one layer of green netting material, which reduced the ambient light intensity by 84% (low light-intensity treatment). The remaining four cages were covered only with one layer of the white netting material, which served as positive control (to control for the effect of netting) and reduced light intensity by 53% (high light-intensity treatment). In other words, the low light-intensity treatment received 66% less light than the high light-intensity treatment.

The experiment totaled 320 pots ([16 no-competition × 2 replicates + 16 intraspecific-competition + 32 interspecific-competition] × 2 nutrient treatments × 2 light treatments). The eight cages were randomly assigned to fixed positions in the botanical garden. The pots were randomly assigned to the eight cages (40 pots in each cage), and were re-randomized once within and across cages of the same light treatment on 3 August 2020. Besides the weekly fertilization, we watered the plants two or three times a week to avoid water limitation. On 7 and 8 September 2020, we harvested the aboveground biomass of all plants. The biomass was dried at 70□ for 96h to constant weight, and then weighed to the nearest 0.1 mg.

### Statistical analyses

All analyses were performed using R version 3.6.1 (R Core Team 2019). To test whether resource availabilities affected competitive outcomes between alien and native species, we applied linear mixed-effects models to analyze the two experiments jointly and separately, using the *nlme* package (Pinheiro *et al*. 2018). For the model used to analyze the two experiments jointly, we excluded interspecific competition between two aliens and between two natives from the experiment in China, because these combinations were not included in the experiment in Germany. When we analyzed each experiment separately, the results were overall similar to the results of the joint analysis. Therefore, we focus in the manuscript on the joint analysis, and present the results of the separate analyses in Supplement S1.

Because plant mortality was low and mainly happened after transplanting, we excluded pots in which plants had died. The final dataset contained 1180 individuals from 871 pots. In the model, we included aboveground biomass of individuals as response variable. We included origin of the species (alien or native), competition treatment (see below for details), nutrient treatment, light treatment and their interactions as fixed effects; and study site (China or Germany), and identity and family of the species as random effects. In addition, we allowed each species to respond differently to the nutrient and light treatments (i.e. we included random slopes). To account for pseudoreplication (Hurlbert 1984), we also included cages as random block effect and pots as random effect. In the competition treatment, we had three levels: 1) no competition, 2) intraspecific competition, and 3) interspecific competition between alien and native species. To split them into two contrasts, we created two dummy variables (Schielzeth 2010) testing 1) the effect of competition, and 2) the difference between intra- and interspecific competition. To improve normality of the residuals, we natural-log-transformed aboveground biomass. To improve homoscedasticity of residuals, we allowed the species and competition treatment to have different variances by using the *varComb* and *varIdent* functions (Zuur *et al*. 2009). Significances of the fixed effects were assessed with likelihood-ratio tests with *car* package(Fox & Weisberg 2018).

To determine the ‘competitive outcome’, i.e. which species will exclude or dominate over the other species at the end point for the community (Gibson *et al*. 1999; Zhang *et al*. 2020), one should ideally conduct a long-term experiment. Alternatively, one could vary the density of each species, which mimics the dynamics of species populations across time (space-for-time-substitution method; see Hart *et al*. 2012; Zhang & van Kleunen 2019 for examples). However, this will largely increase the size of the experiment, especially when combining with other treatments. Still, by growing plants grown alone, in intraspecific competition and in interspecific competition, our experiments meet the minimal requirement for measuring competitive outcome (Hart *et al*. 2018; Zhang *et al*. 2020). In the linear mixed-effects model of individual biomass, a significant effect of origin would indicate that alien and native species differed in their biomass production, across all competition and resource-availability (light and nutrients) treatments. This would tell us the competitive outcome between aliens and natives across different resource availabilities. For example, an overall higher level of biomass production of alien species would indicate that aliens would dominate when competing with natives. A significant interaction between resource-availability treatment and origin of the species would indicate that resource availabilities affect the biomass production of alien and native species differently, averaged across all competition treatments. In other words, it would indicate that resource availabilities affect the competitive outcome between aliens and natives.

A significant interaction between a resource-availability treatment and the competition treatment would indicate that resource availabilities affect the effect of competition (e.g. no competition *vs*. competition). Note that, most studies infer competitive outcome from effect of competition, for example, by calculating relative interaction intensity (Armas *et al*. 2004). However, while competitive outcome and effect of competition are often related, they are not equivalent (Gibson *et al*. 1999). This is because competitive outcome is both determined by effect of competition and intrinsic growth rate (Hart *et al*. 2018; Zhang & van Kleunen 2019). For example, a plant species that strongly suppresses other species but has a low intrinsic growth rate still cannot dominate the community.

## Results

Overall, aboveground biomass production of plants increased with nutrient availability (+66.6%; Fig. 2; Table S2; χ^2^ = 46.42, *P* < 0.001), and marginally increased with light intensity (+67.9%; χ^2^ = 3.18, *P* = 0.075). Moreover, aboveground biomass production increased the most with a joint increase of nutrient availability and light intensity (+79.4%), as indicated by the interaction between nutrient and light treatments (χ^2^ = 24.12, *P* < 0.001). Although alien species tended to produce more aboveground biomass than native ones, averaged across competition treatments and the different light and nutrient treatments, this difference was not significant (χ^2^ = 2.04, *P* = 0.153). This indicates that, on average, aliens did not outcompete natives. However, the competitive outcome between aliens and natives depended on the combination of nutrient and light treatments (Fig. 2; Table S2; χ^2^ = 4.66, *P* = 0.031). More specifically, with a joint increase of nutrients and light intensity, aliens produced more (+110.8%) aboveground biomass than natives; whereas this difference was much smaller under low resource availabilities (+48.3%) or with addition of only one type of resource (+48.4% under high nutrients only; +68.9% under high light intensity only).

**Figure 2.**
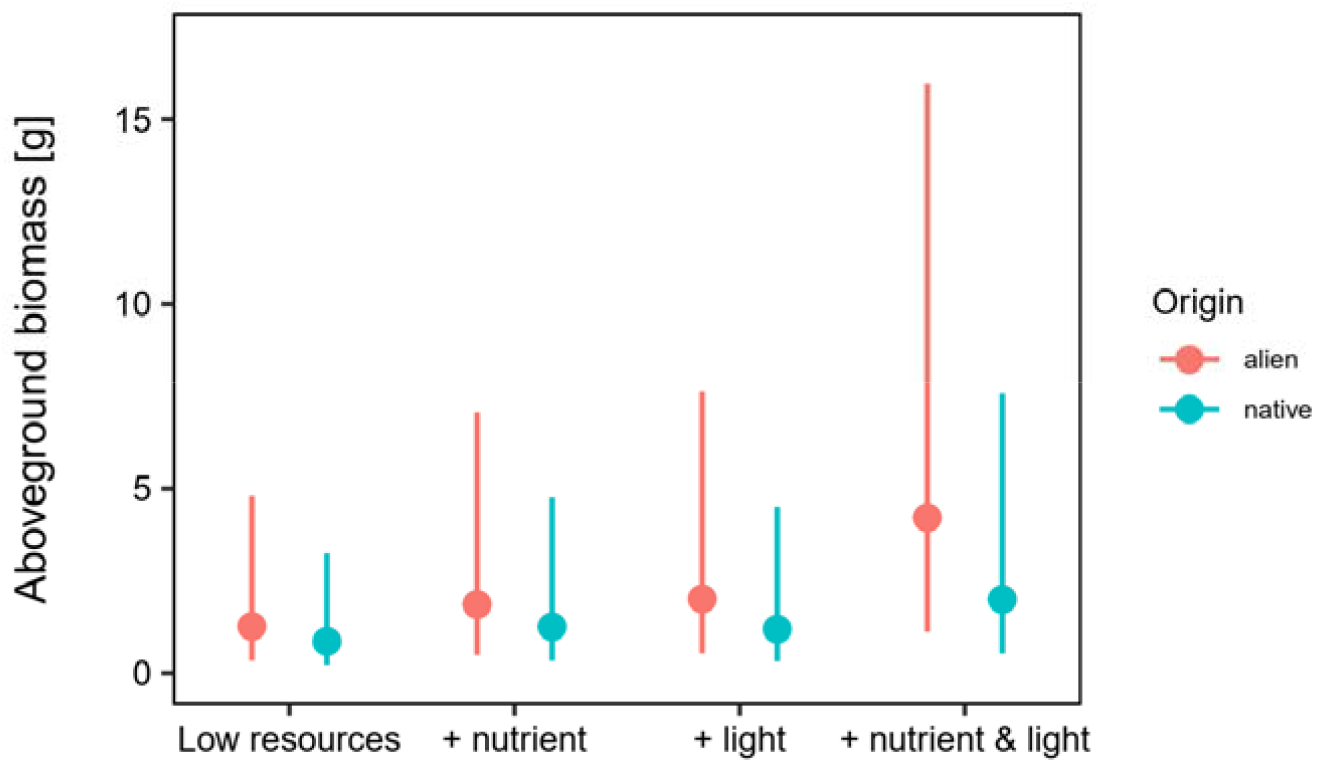
Effects of nutrient and light availabilities on competitive outcomes between alien (red) and native (blue) plants. Competitive outcome is indicated by the difference in average biomass production between alien and native plants across competition treatments (i.e. without competition, intra- and interspecific competition). For example, a higher biomass production of alien plants indicates that aliens outcompete natives. Error bars indicate 95% CIs. The effects of nutrient and light availabilities on the difference in aboveground production between alien and native plants did not significantly depend on competition treatments (see Fig. S4 for details).

Competition reduced (−26.0%) aboveground biomass production, as indicated by the difference between plants grown without competition and plants grown with competition (Fig. 3; Table S2; χ^2^ = 1.93, *P* = 0.165). Although this effect of competition was not statistically significant averaged across the different nutrient or light treatments, it became more apparent with increased nutrients (χ^2^ = 5.21, *P* = 0.022) and increased light intensity (χ^2^ = 4.21, *P* = 0.040). In addition, we found that plants produced more (+16.4%) aboveground biomass when competing with interspecific competitors than with intraspecific competitors (Fig. 3; Table S2; χ^2^ = 20.05, *P* < 0.001).

**Figure 3.**
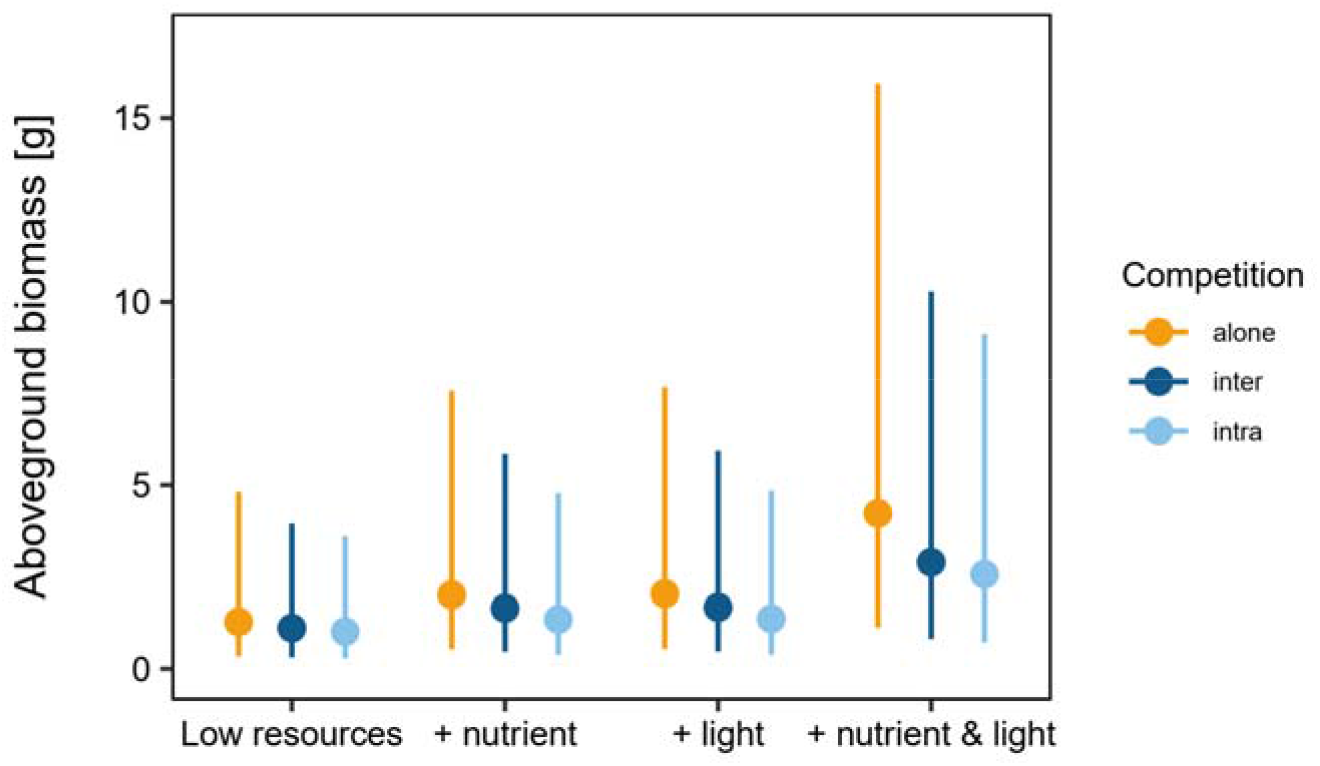
Effects of nutrient and light availabilities on effects of intra- and interspecific competition on plant aboveground biomass. Yellow, dark blue and light blue lines represent plants without competition, and with inter- and intraspecific competition, respectively. Error bars indicate 95% CIs.

## Discussion

We found that under low resource availabilities, alien and native plants did not differ in biomass production, indicating that under those conditions aliens will not outcompete natives. Although an increase in one type of resource, either nutrients or light, increased biomass production, it affected aliens and natives similarly, and thus did not change the potential competitive outcome. However, with a joint increase of nutrients and light intensity, aliens produced more biomass than natives, indicating that aliens will outcompete natives under high availabilities of both resources. Our finding thus supports the fluctuating resource hypothesis, which predicts that ‘a plant community becomes more susceptible to invasion whenever there is an increase in the amount of unused resources’. Furthermore, our finding, along with those of others (Seabloom *et al*. 2015; Liu *et al*. 2016), explains why plant invasion is frequently associated with disturbance. This is because disturbance could result in a flush of nutrient availability and create open patches with a higher light intensity (Wilson & Tilman 1993), a combination that favors naturalized alien plants.

Our finding that across two experiments, addition of one type of resource did not favor alien plants has several implications. First, it suggests that plants —irrespectively of their origin— are limited by multiple factors, such as nutrients, light, water and herbivory. In other words, niche space has multiple dimensions, each of which is represented by one limiting factor. While addition of one resource removes one dimension from the niche space, the remaining dimensions could still limit both alien and native plants, maintaining coexistence of aliens and natives. Some previous studies, in line with our finding, showed that addition of one type of resource (nutrients) did not favor alien plants (Liu *et al*. 2018). However, others found that addition of nutrients only was sufficient to favor alien plants (Parepa *et al*. 2013; Liu & van Kleunen 2017). One explanation for the apparent discrepancy could be that the latter studies were conducted under high light conditions. This was likely the case as the latter two studies were done in summer, while Liu *et al*. (2018) was done in a greenhouse in winter. With addition of nutrients, their environments were similar to joint increases of nutrients and light in our study, which reduced more dimensions of the niche space, favoring dominance by one of the two species.

A second implication is that our finding suggests that alien and native species did not strongly differ in their competitive abilities for nutrients and light. As addition of one resource removes one dimension from the niche space, it intensifies competition for other dimensions. For example, nutrient addition can intensify light competition (Hautier *et al*. 2009), which is also indicated by our finding that competition was more severe with nutrient addition (Fig. 3a). Consequently, if alien plants have stronger competitive abilities for light (e.g. have a lower minimum requirement of light) than natives, they will dominate with nutrient addition. However, as this was not the case, we conclude that there was no strong difference in competitive abilities for light between the aliens and natives.

The two implications mentioned above raise the question which factor or factors determine the higher competitiveness of naturalized alien plants with joint increases of nutrients and light. One potential factor could be plant enemies, as we indeed did observe herbivory in our experiments. Because the evolutionary histories of alien plants differ from those of the native plants, alien plants might be released from natural enemies (Keane & Crawley 2002). This advantage might be stronger when other factors, for example, resource availabilities, are not limiting the plants (Coley *et al*. 1985; Blumenthal 2006). An alternative potential factor is preadaptation of naturalized alien plants. Many alien plants occur in human-associated habitats (Kalusová *et al*. 2017), where resource availabilities are high due to human disturbance. Consequently, of the many alien plants that have been introduced, the ones that managed to naturalize or become invasive are most likely the ones that were selected for high growth rates under high resource availabilities. Given that these two explanations are not mutually exclusive, future studies that test their relative importance are needed.

## Conclusions

The fluctuating resource hypothesis suggests that, with an increase in resources, a plant community becomes more susceptible to invasion. Our study suggests that this is particularly the case with increases of multiple resources, as this could greatly reduce the dimensionality of niche space, leading to competitive exclusion of one of the species. This can also explain why many studies have found that biological invasions are more frequent in disturbed, high resource environments.

## Acknowledgements

We thank O. Ficht, M. Fuchs, X. Zhang, L. Wang and Y. Li for practical assistance. ZZ acknowledges funding from the China Scholarship Council (201606100049) and support from the International Max Planck Research School for Organismal Biology. YL acknowledges funding from the Chinese Academy of Sciences (Y9B7041001).

## Notes

### Competing Interest Statement

The authors have declared no competing interest.

