## Supplement for "Increases in multiple resources promote plant invasion"

Supporting information

### Supplement S1 The species used in the experiments

**Table S1** The 24 species used in the experiment in China and the experiment in Germany.

| **Species** | **Site** | **Family** | **Origin** | **Propagule** | **Sowing date** | **Propagule source** |
| --- | --- | --- | --- | --- | --- | --- |
| *Alternanthera philoxeroides* | China | Amaranthaceae | alien | stem fragment | 24 June 2020 | wild population |
| *Alternanthera sessilis* | China | Amaranthaceae | native | stem fragment | 23 June 2020 | wild population |
| *Bidens pilosa* | China | Compositae | alien | seed | 27 June 2020 | wild population |
| *Solidago canadensis* | China | Compositae | alien | seed | 21 May 2020 | wild population |
| *Bidens maximowicziana* | China | Compositae | native | seed | 27 June 2020 | wild population |
| *Solidago decurrens* | China | Compositae | native | seed | 21 May 2020 | wild population |
| *Paspalum notatum* | China | Poaceae | alien | seed | 24 June 2020 | commercial |
| *Paspalum orbiculare* | China | Poaceae | native | stem fragment | 24 June 2020 | wild population |
| *Diplotaxis muralis* | Germany | Brassicaceae | alien | seed | 15 June 2020 | Boga Uni-Konstanz^1^ |
| *Cardamine hirsuta* | Germany | Brassicaceae | native | seed | 15 June 2020 | Boga Uni-Konstanz |
| *Galinsoga parviflora* | Germany | Compositae | alien | seed | 15 June 2020 | commercial |
| *Hieracium pilosella* | Germany | Compositae | native | seed | 15 June 2020 | commercial |
| *Elsholtzia cilliata* | Germany | Lamiaceae | alien | seed | 15 June 2020 | Boga Uni-Konstanz |
| *Salvia verticilliata* | Germany | Lamiaceae | alien | seed | 15 June 2020 | commercial |
| *Mentha longifolia* | Germany | Lamiaceae | native | seed | 15 June 2020 | commercial |
| *Prunella vulgaris* | Germany | Lamiaceae | native | seed | 15 June 2020 | commercial |
| *Onobrychis viciifolia* | Germany | Leguminosae | alien | seed | 15 June 2020 | commercial |
| *Vicia villosa* | Germany | Leguminosae | alien | seed | 15 June 2020 | commercial |
| *Medicago falcata* | Germany | Leguminosae | native | seed | 15 June 2020 | commercial |
| *Medicago lupulina* | Germany | Leguminosae | native | seed | 15 June 2020 | commercial |
| *Epilobium ciliatum* | Germany | Onagraceae | alien | seed | 15 June 2020 | Boga Uni-Konstanz |
| *Epilobium hirsutum* | Germany | Onagraceae | native | seed | 15 June 2020 | Boga Uni-Konstanz |
| *Eragrostis minor* | Germany | Poaceae | alien | seed | 15 June 2020 | commercial |
| *Vulpia myuros* | Germany | Poaceae | native | seed | 15 June 2020 | commercial |

^1^Botanical Garden of the University of Konstanz

### Supplement S2 The separate analyses on the two experiments

#### Statistical analyses

##### The experiment in China

The statistical model was similar to the one that we used in the joint analysis of both experiments together. In brief, we included aboveground biomass as response variable; origin of the species (alien or native), competition treatment, nutrient treatment, light treatment and their interactions as fixed effects; and identity and family of the species as random effects. In addition, we allowed each species to respond differently to the nutrient and light treatments (i.e. we included random slopes). However, in this model, the competition treatment had four instead of three levels: 1) competition-free, 2) intraspecific competition, 3) interspecific competition from alien species, and 4) interspecific competition from native species. We created three dummy variables to split the competition treatment into three contrasts to test 1) the effect of competition, 2) the difference between intra- and inter-specific competition, and 3) the difference between interspecific competition from alien species and that from native species.

##### The experiment in Germany

The model was the same as for the experiment in China. However, like in the joint analysis, the competition treatment had only three levels: 1) competition-free, 2) intraspecific competition, and 3) interspecific competition between alien and native species. Therefore, we used the same dummy variables as in the joint analysis.

#### Results

Consistent with the joint analysis, both experiments showed that biomass production of plants increased with increases of nutrients and light intensity (Fig. S1a&b; Table S2). In addition, both experiments found that the competitive outcome between aliens and natives was affected by the interaction between nutrient and light treatments (Fig. S1a&b; Table S2). More specifically, under low resource availabilities or with addition of only one type of resources, biomass production did not differ between aliens and natives, whereas with a joint increase of nutrients and light, aliens produced more biomass than natives.

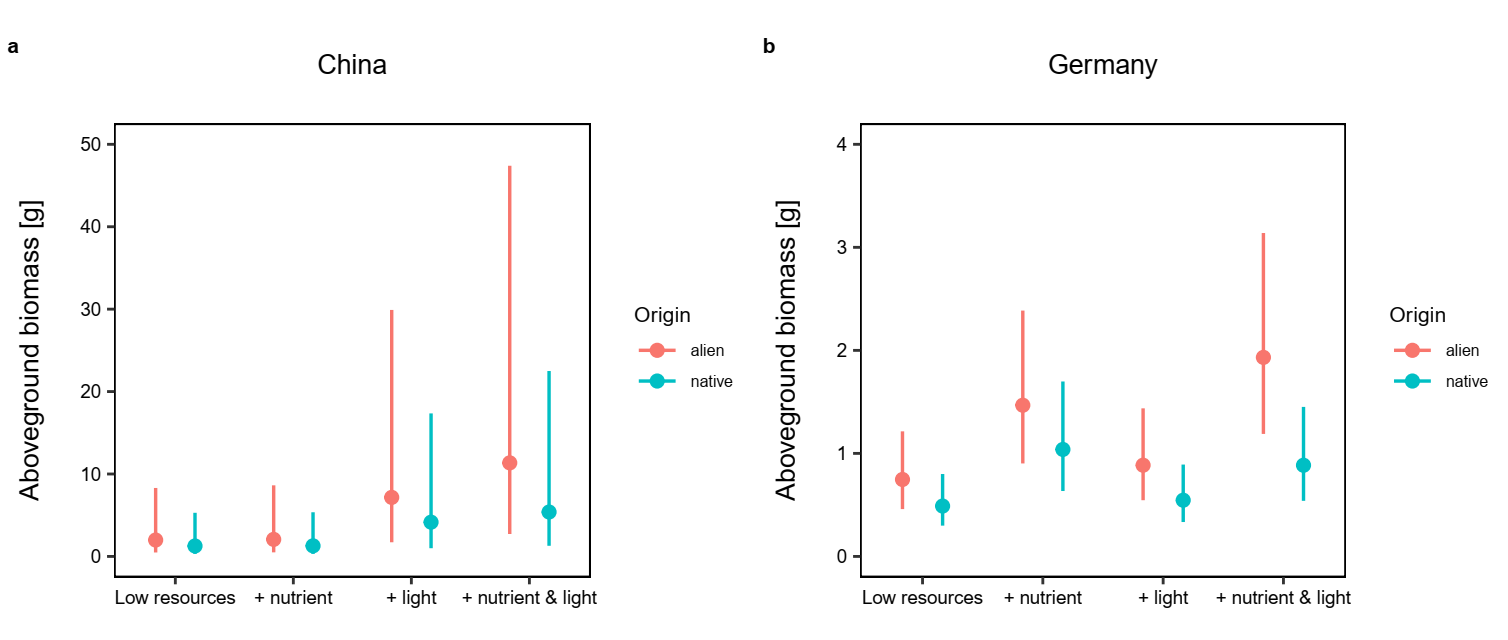

**Figure S1** Effects of nutrient and light availabilities on competitive outcomes between alien (red) and native (blue) plants. Data from the experiments in China (a) and Germany (b) were analyzed separately Competitive outcome is indicated by the difference in average biomass production. For example, a higher biomass production of alien plants indicates that aliens outcompete natives. Error bars indicate 95% CIs.

Consistent with the joint analysis, the experiment in China showed that biomass production largely increased with a joint increase of nutrients and light, as indicated by the interaction between nutrient and light treatments (Fig. S1a; Table S2). Furthermore, this experiment included competition between two aliens and that between two natives. It showed that the origin of competitor species also matters (Fig. S2): aliens produced more biomass when competing with natives than when competing with other aliens, whereas biomass production of natives was not strongly affected by the origin of competitor species.

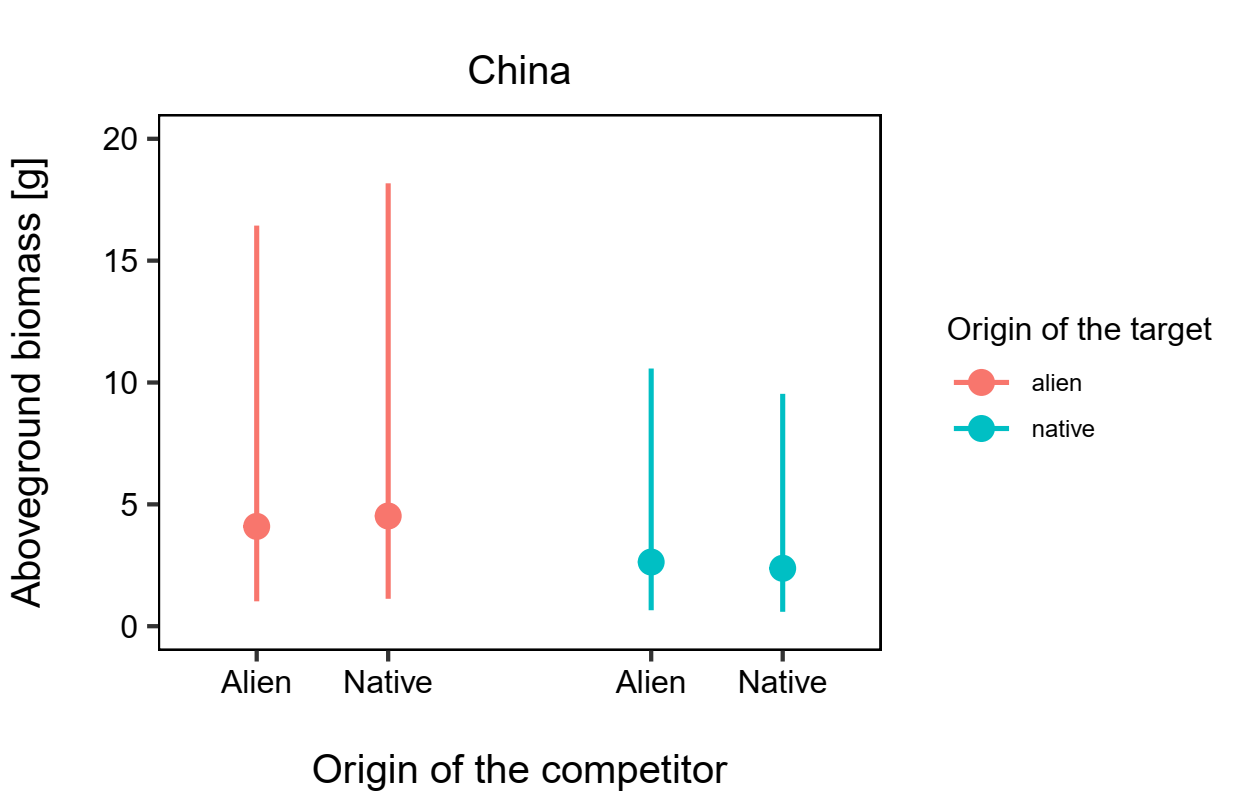

**Figure S2** Competitive outcomes among alien and native plants for the experiment in China. Red and blue colors indicate alien and native species, respectively, which were grown with interspecific competitors. The competitors were either an alien species or a native. Error bars indicate 95% CIs.

The experiment in Germany showed that the competitive outcome between alien and native species was affected by light intensity (Fig. S1b). Under low light intensity, biomass production did not strongly differ between aliens and natives, whereas aliens produced more biomass than natives with an increase of light intensity. This effect is mainly driven by the fact that aliens produced more biomass than natives with a joint increase of nutrients and light intensity (Fig. S1b), and by the fact that aliens produced more biomass than native under interspecific competition and high light-intensity (Fig. S3)

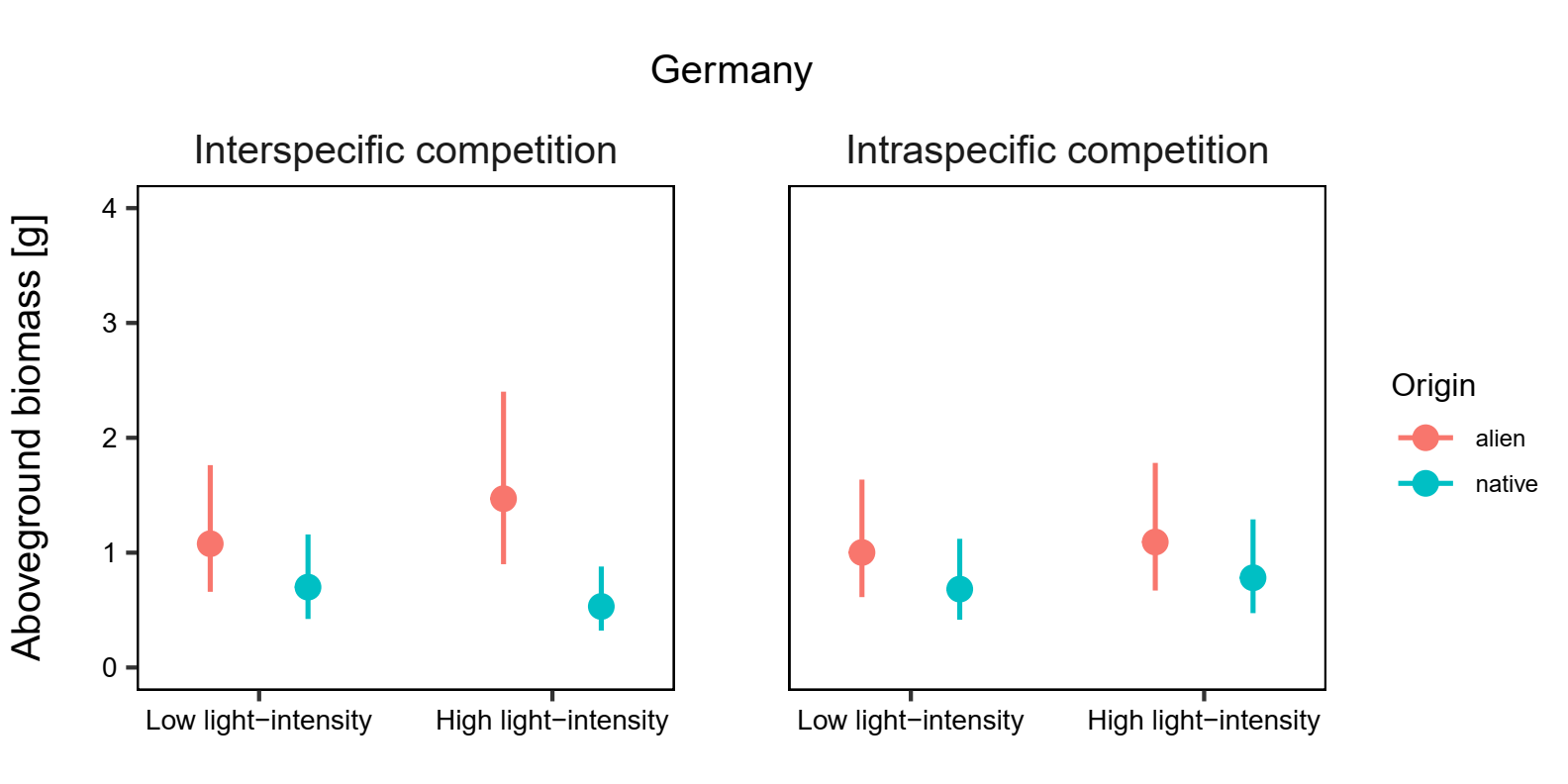

**Figure S3** Effect of light availability on alien (red) and native (blue) plants under inter- or intraspecific competition. Error bars indicate 95% CIs.

**Table S2** Effect of origin of target species, competition treatments and resource availabilities on aboveground biomass of plants.

|  |  | | China | |  | |  |  | Germany |  |  |  | Joint |  |  |
| --- | --- | --- | --- | --- | --- | --- | --- | --- | --- | --- | --- | --- | --- | --- | --- |
| Items | DF | χ^2^ | | p-value | |  | | DF | χ^2^ | p-value |  | DF | χ^2^ | p-value |  |
| origin | 1 | 0.301 | | 0.583 | |  | | 1 | 3.637 | 0.057 | † | 1 | 2.042 | 0.153 |  |
| T_alone_comp^1^ | 1 | 3.313 | | 0.069 | | † | | 1 | 1.976 | 0.16 |  | 1 | 2.087 | 0.149 |  |
| T_intra_inter^1^ | 1 | 20.281 | | <0.001 | | * | | 1 | 0.199 | 0.655 |  | 1 | 20.054 | <0.001 | * |
| T_alien_native^2^ | 1 | 0.022 | | 0.881 | |  | | - | - | - |  | - | - | - |  |
| fert | 1 | 5.55 | | 0.018 | | * | | 1 | 99.935 | <0.001 | * | 1 | 46.424 | <0.001 | * |
| light | 1 | 44.059 | | <0.001 | | * | | 1 | 5.626 | 0.018 | * | 1 | 3.176 | 0.075 | † |
| origin:T_alone_comp | 1 | 0.146 | | 0.702 | |  | | 1 | 1.152 | 0.283 |  | 1 | 0.626 | 0.429 |  |
| origin:T_intra_inter | 1 | 0.833 | | 0.362 | |  | | 1 | 2.588 | 0.108 | † | 1 | 0.355 | 0.551 |  |
| origin:T_alien_native | 1 | 8.761 | | 0.003 | | * | | - | - | - |  | - | - | - |  |
| origin:fert | 1 | 0.434 | | 0.51 | |  | | 1 | 0.442 | 0.506 |  | 1 | 0.817 | 0.366 |  |
| T_alone_comp:fert | 1 | 0.188 | | 0.665 | |  | | 1 | 21.74 | <0.001 | * | 1 | 5.209 | 0.022 | * |
| T_intra_inter:fert | 1 | 0.285 | | 0.594 | |  | | 1 | 0.86 | 0.354 |  | 1 | 0.002 | 0.96 |  |
| T_alien_native:fert | 1 | 0.141 | | 0.707 | |  | | - | - | - |  | - | - | - |  |
| origin:light | 1 | 0.183 | | 0.669 | |  | | 1 | 4.698 | 0.03 | * | 1 | 1.602 | 0.206 |  |
| T_alone_comp:light | 1 | 2.718 | | 0.099 | |  | | 1 | 1.674 | 0.196 |  | 1 | 4.214 | 0.040 | * |
| T_intra_inter:light | 1 | 0.107 | | 0.744 | |  | | 1 | 0.207 | 0.649 |  | 1 | 0.021 | 0.884 |  |
| T_alien_native:light | 1 | <0.001 | | 0.995 | |  | | - | - | - |  | - | - | - |  |
| fert:light | 1 | 43.581 | | <0.001 | | * | | 1 | 0.026 | 0.872 |  | 1 | 24.122 | <0.001 | * |
| origin:T_alone_comp:fert | 1 | 0.409 | | 0.523 | |  | | 1 | 0.013 | 0.911 |  | 1 | 1.91 | 0.167 |  |
| origin:T_intra_inter:fert | 1 | 0.845 | | 0.358 | |  | | 1 | 1.123 | 0.289 |  | 1 | 0.011 | 0.918 |  |
| origin:T_alien_native:fert | 1 | 0.205 | | 0.65 | |  | | - | - | - |  | - | - | - |  |
| origin:T_alone_comp:light | 1 | 0.245 | | 0.621 | |  | | 1 | 0.28 | 0.596 |  | 1 | 0.061 | 0.805 |  |
| origin:T_intra_inter:light | 1 | 0.681 | | 0.409 | |  | | 1 | 7.099 | 0.008 | * | 1 | 3.065 | 0.080 | † |
| origin:T_alien_native:light | 1 | 0.419 | | 0.518 | |  | | - | - | - |  | - | - | - |  |
| origin:fert:light | 1 | 3.435 | | 0.064 | | † | | 1 | 3.839 | 0.050 | * | 1 | 4.655 | 0.031 | * |

continued:

|  |  | China |  |  |  | Germany |  |  |  | Pooled |  |
| --- | --- | --- | --- | --- | --- | --- | --- | --- | --- | --- | --- |
| items | DF | χ^2^ | p-value |  | DF | χ^2^ | p-value |  | DF | χ^2^ | p-value |
| T_alone_comp:fert:light | 1 | 0.566 | 0.452 |  | 1 | 0.906 | 0.341 |  | 1 | 0.021 | 0.884 |
| T_intra_inter:fert:light | 1 | 3.003 | 0.083 |  | 1 | 1.421 | 0.233 |  | 1 | 2.412 | 0.120 |
| T_alien_native:fert:light | 1 | 0.759 | 0.384 |  | - | - | - |  | - | - | - |
| origin:T_alone_comp:fert:light | 1 | 0.168 | 0.682 |  | 1 | 3.237 | 0.072 | † | 1 | 0.871 | 0.351 |
| origin:T_intra_inter:fert:light | 1 | 0.313 | 0.576 |  | 1 | 1.611 | 0.204 |  | 1 | 0.932 | 0.334 |
| origin:T_alien_native:fert:light | 1 | 1.627 | 0.202 |  | - | - | - |  | - | - | - |
| **Random effects** | SD |  |  |  | SD |  |  |  | SD |  |  |
| Study site | - |  |  |  | - |  |  |  | 0.843 |  |  |
| Family | 0.007 |  |  |  | 0.286 |  |  |  | 0.178 |  |  |
| Species | 1.371 |  |  |  | 0.509 |  |  |  | 0.844 |  |  |
| Family of the competitor | 0.002 |  |  |  | 0.112 |  |  |  | 0.001 |  |  |
| Competitor species | 0.153 |  |  |  | 0.176 |  |  |  | 0.204 |  |  |
| light:species | 0.417 |  |  |  | 0.070 |  |  |  | 0.302 |  |  |
| fert:species | 0.155 |  |  |  | 0.125 |  |  |  | 0.217 |  |  |
| cage_first^3^ | - |  |  |  | <0.001 |  |  |  | 0.284 |  |  |
| cage_second^3^ | - |  |  |  | <0.001 |  |  |  | <0.001 |  |  |
| pot | <0.001 |  |  |  | <0.001 |  |  |  | <0.001 |  |  |
| Residual | 0.374 |  |  |  | 0.726 |  |  |  | 0.346 |  |  |

^1^The competition treatments were split into contrasts to test the effect of competition (T_alone_comp) and the difference between intra- and interspecific competition (T_intra_inter).

^2^For the experiment of China, which included competition between two natives and that between two aliens, there was an additional contrast to test the difference between interspecific competition from aliens and that from natives (T_alien_native).

^3^For the experiment of Germany, pots were randomized twice across cages, so that there were two ‘cage’ (i.e. block) effect.

### Supplement S3 Effects of nutrient and light availabilities

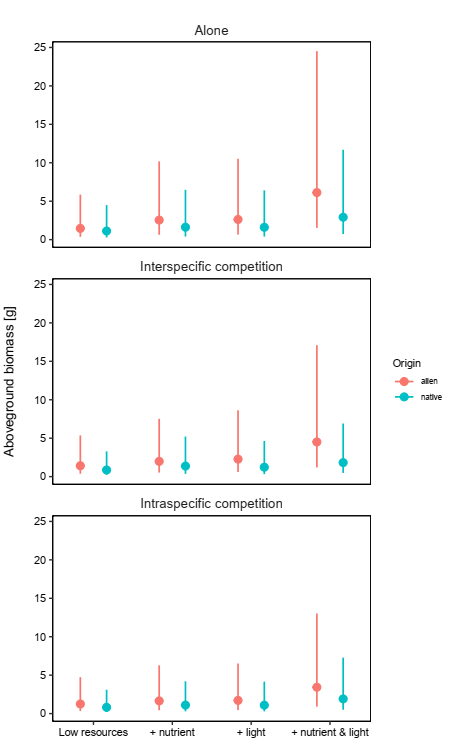

**Figure S4** Effects of nutrient and light availabilities on aboveground biomass of alien (red) and native (blue) plants. The plants without competition (upper), with inter- (middle) and intraspecific competition are shown separately. Similar with Fig. 2, aliens had higher aboveground biomass than natives with addition of both nutrients and light. This effect did not significantly depend on competition treatment (i.e. no significant effect of origin:T_alone_comp:fert:light, origin:T_intra_inter:fert:light or origin:T_alien_native:fert:light in **Table S2**).
